# Deciphering Cancer Genomes with GenomeSpy: A Grammar-Based Visualization Toolkit

**DOI:** 10.1101/2023.10.06.561159

**Authors:** Kari Lavikka, Jaana Oikkonen, Yilin Li, Taru Muranen, Giulia Micoli, Giovanni Marchi, Alexandra Lahtinen, Kaisa Huhtinen, Rainer Lehtonen, Sakari Hietanen, Johanna Hynninen, Anni Virtanen, Sampsa Hautaniemi

## Abstract

**Background:** Visualization is an indispensable facet of genomic data analysis. Despite the abundance of specialized visualization tools, there remains a distinct need for tailored solutions. However, their implementation typically requires extensive programming expertise from bioinformaticians and software developers, especially when building interactive applications. Toolkits based on visualization grammars offer a more accessible, declarative way to author new visualizations. Nevertheless, current grammar-based solutions fall short in adequately supporting the interactive analysis of large data sets with extensive sample collections, a pivotal task often encountered in cancer research.

**Results:** We present GenomeSpy, a grammar-based toolkit for authoring tailored, interactive visualizations for genomic data analysis. Users can implement new visualization designs with little effort by using combinatorial building blocks that are put together with a declarative language. These fully customizable visualizations can be embedded in web pages or end-user-oriented applications. The toolkit also includes a fully customizable but user-friendly application for analyzing sample collections, which may comprise genomic and clinical data. Findings can be bookmarked and shared as links that incorporate provenance information. A distinctive element of GenomeSpy’s architecture is its effective use of the graphics processing unit (GPU) in all rendering. GPU usage enables a high frame rate and smoothly animated interactions, such as navigation within a genome. We demonstrate the utility of GenomeSpy by characterizing the genomic landscape of 753 ovarian cancer samples from patients in the DECIDER clinical trial. Our results expand the understanding of the genomic architecture in ovarian cancer, particularly the diversity of chromosomal instability. We also show how GenomeSpy enabled the discovery of clinically actionable genomic aberrations.

**Conclusions:** GenomeSpy is a visualization toolkit applicable to a wide range of tasks pertinent to genome analysis. It offers high flexibility and exceptional performance in interactive analysis. The toolkit is open source with an MIT license, implemented in JavaScript, and available at https://genomespy.app/.

## Background

Effective visualization facilitates hypothesis generation and the assessment of automatic analyses, making it an indispensable facet of genomic data analysis [1]. However, interpreting complex genomic data sets calls for visualization methods tailored to the analyzed data [2], a need underscored by the availability of numerous special-purpose tools [3, 4]. Implementing tailored visualizations, particularly those that offer interactivity, typically necessitates developing new software packages from scratch or writing plugins for existing ones, such as the modular JBrowse 2 [5] genome browser. This laborious process demands considerable programming expertise that is beyond the scope of most bioinformaticians.

Visualization grammars like ggplot2 [6], Vega-Lite [7], and the genomic-data-focused Gosling [8] and ggbio [9], which all build upon the concept initially presented in the Grammar of Graphics [10], support tailored visualizations with a more accessible approach: instead of using an imperative programming language, they are specified using combinatorial building blocks such as graphical marks, scales, transformations, and view compositions, which are put together using a declarative language. However, none of these grammar-based solutions sufficiently cater to the typical analysis task in cancer research: the exploration and analysis of large sample collections to find patterns and outliers in cohorts. They either lack support for genomic data, fail to visualize numerous concurrent samples, disallow interactive filtering and grouping, or underperform with large data sets.

Herein, we present GenomeSpy, a toolkit designed to simplify the crafting of interactive visualizations and empower end users to effectively explore and analyze large data sets, particularly in cancer research. The toolkit features a grammar that enables effortless implementation of different visualization strategies (Figure 1). This characteristic makes GenomeSpy fundamentally distinct from genome browsers, such as IGV [11], igv.js [12], JBrowse 2, and UCSC Genome Browser [13], which comprise pre-defined track types designed for specific data formats that are displayed using rigid visual encodings. In addition, we incorporated the grammar into an analysis application for sample collections, with a pronounced focus on fluid interaction. This design principle aims to make interaction with visualizations more rewarding, ultimately enhancing users’ performance [14]. Fluid interaction changes browsing and exploration, which are considered a rate-limiting step in data analysis [2], into an endeavor that fosters insights.

**Figure 1:**
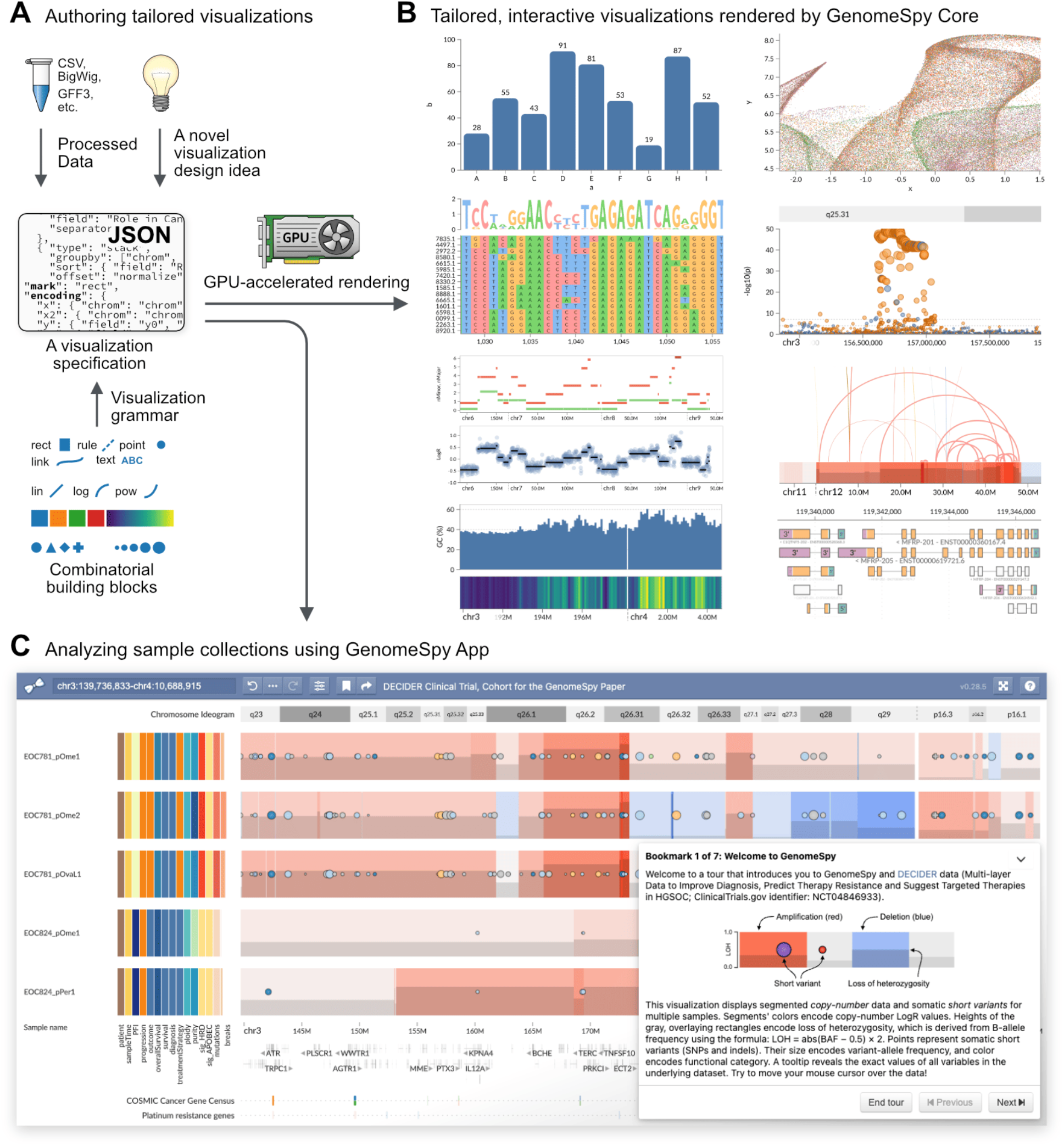
Overview of GenomeSpy. **A** GenomeSpy enables tailored visualizations through its JSON-based visualization grammar, which defines how the building blocks, such as marks and scales, can be combined into a visualization specification. Instead of relying on pre-defined templates or track types, the user can freely compose visualizations from various graphical marks and map data attributes to different visual channels, such as color and position. **B** GenomeSpy core library parses the specification and renders it using GPU-accelerated graphics to ensure smooth interactions such as zooming and panning. Interactive versions of the above examples are available at https://genomespy.app/. **C** GenomeSpy App builds upon the core and enables the analysis of sample collections. The above visualization with 753 samples is available for exploration at https://csbi.ltdk.helsinki.fi/p/genomespy-preprint/

We demonstrate the utility and key features of GenomeSpy by exploring and analyzing 753 whole-genome-sequenced (WGS) samples from 215 patients who belong to prospective, longitudinal, multi-region observational study DECIDER (Multi-layer Data to Improve Diagnosis, Predict Therapy Resistance and Suggest Targeted Therapies in HGSOC; ClinicalTrials.gov identifier: NCT04846933) that started recruitment in 2012. The DECIDER trial focuses on characterizing and overcoming therapy resistance in ovarian high-grade serous carcinoma (HGSC), the most common and aggressive epithelial ovarian cancer subtype. The standard-of-care (SOC) for HGSC consists of debulking surgery and platinum-taxane chemotherapy, often combined with maintenance therapy with ADP ribose polymerase (PARP) or VEGF pathway inhibitors [15]. While ∼80% of HGSC patients respond well to the SOC, most of the patients suffer from recurrence and rapid disease progression leading to five-year survival rate of only <40% [16]. Except for nearly 100% prevalent TP53 mutations, HGSC lacks recurrent mutations but is characterized by complex genomes with large-scale copy-number alterations, hindering a deeper mechanistic understanding of the disease [17, 18]. Furthermore, diagnosis is often complicated by rare morphologic and molecular traits [19, 20]. Herein, our hypothesis is that interpreting large genomics data sets from genomically complex cancers, such as HGSC, requires tailored visualization methods, such as one built with GenomeSpy.

## Results

GenomeSpy is a JavaScript-based toolkit that allows developers and bioinformaticians to build interactive visualizations for genome analysis. To construct such a visualization, a user writes a visualization specification in JavaScript Object Notation (JSON) format, adhering to the rules of the visualization grammar (Figure 1A). GenomeSpy’s grammar draws inspiration from the design principles of Vega-Lite, a high-level grammar of interactive graphics [7], enhancing it for robust support of genomic data (Supplementary Note). Figure 2 demonstrates GenomeSpy’s grammar-based approach with a typical use case: a nucleotide sequence of a reference genome.

**Figure 2:**
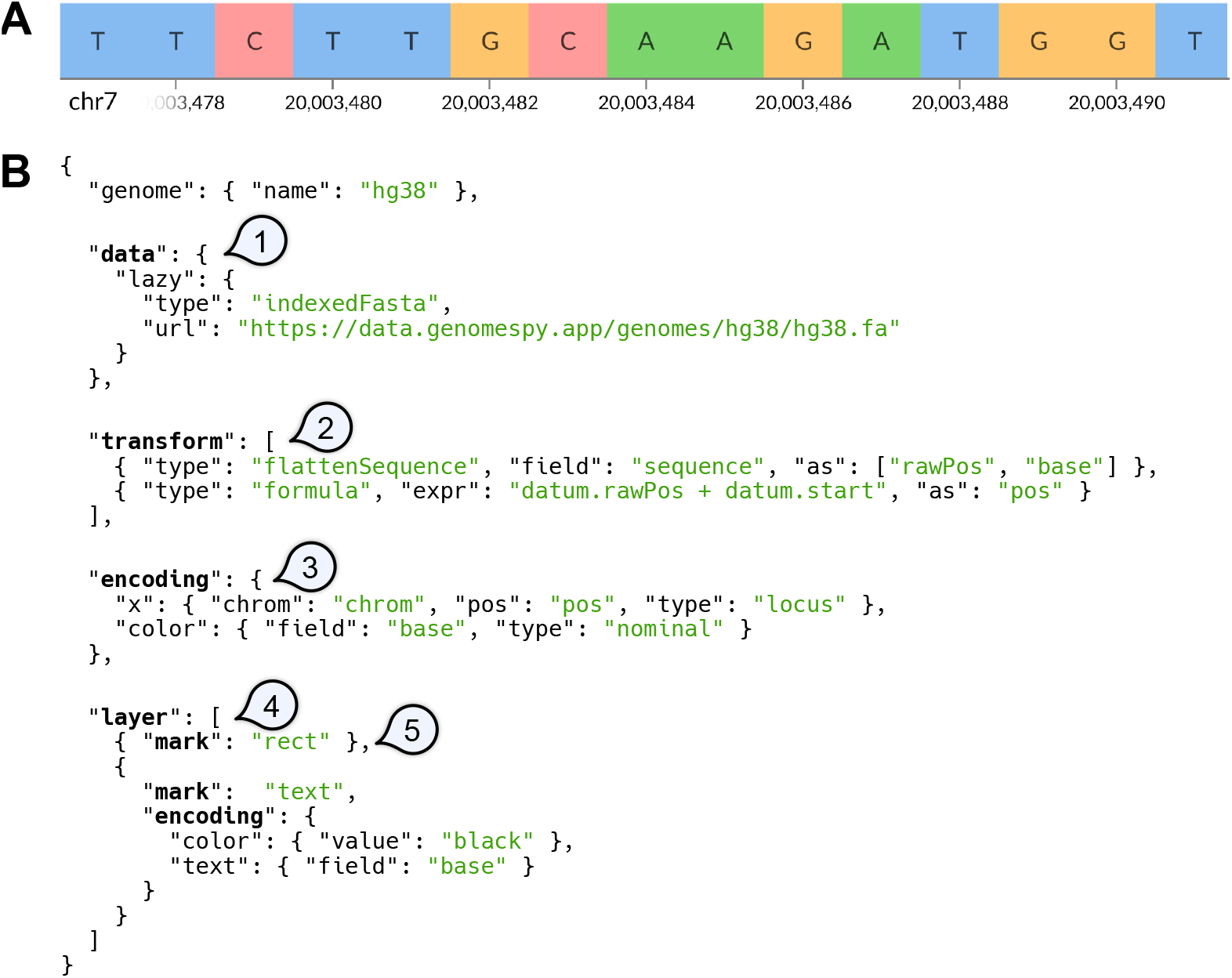
Specifying a visualization of a reference nucleotide sequence using the grammar. **A** The example visualization comprises letters that are superimposed on colored rectangles. The genomic axis is generated automatically. **B** The GenomeSpy core library provides no predefined track types. Instead, the visualization author supplies a JSON-based specification that defines how the building blocks are put together. (1) The *data* property specifies a data source. In this example, data are loaded lazily from an indexed FASTA file as the user navigates the genome. (2) Optional *transformations* modify the data stream. Here, the sequence strings provided by the data source are split into data objects representing individual nucleotides with their coordinates. (3) The *encoding* property allows mapping data fields to visual channels. The x axis is treated as genomic coordinates, as it has a “locus” data type. (4) The *layer* property composes multiple child views by layering them. (5) The *mark* property specifies the graphical mark to be used in a view. Here, “rect” is used for the background rectangles and “text” for the bases. N.B. The specification has been simplified for clarity by omitting non-critical properties. A complete example is available in GenomeSpy’s documentation.

The *core library* constitutes the toolkit’s main component. It implements the grammar and renders the visualization according to the provided specification (Figure 1B). The library can serve as a component in JavaScript web applications, or it can be embedded on web pages such as Observable notebooks (https://observablehq.com/collection/@tuner/genomespy). An example of a special-purpose application built using the core library is SegmentModel Spy (Figure 3, Supplementary Note), which allows a comprehensive assessment of copy-number segmentation output from the Genome Analysis Toolkit (GATK) [21]. A crucial element in the core library’s architecture is its use of GPU acceleration through the WebGL 2 API for all graphics and scale transformations (Supplementary Note). GPU acceleration enables efficient rendering with a high frame rate and minimal latency, which facilitates insight generation [22]. It also allows fluid, smoothly animated interactions, such as continuous zooming and panning in large data sets. While smooth transition animations make the user experience attractive, they have also been shown to improve users’ perception of causality during interactions [23].

**Figure 3:**
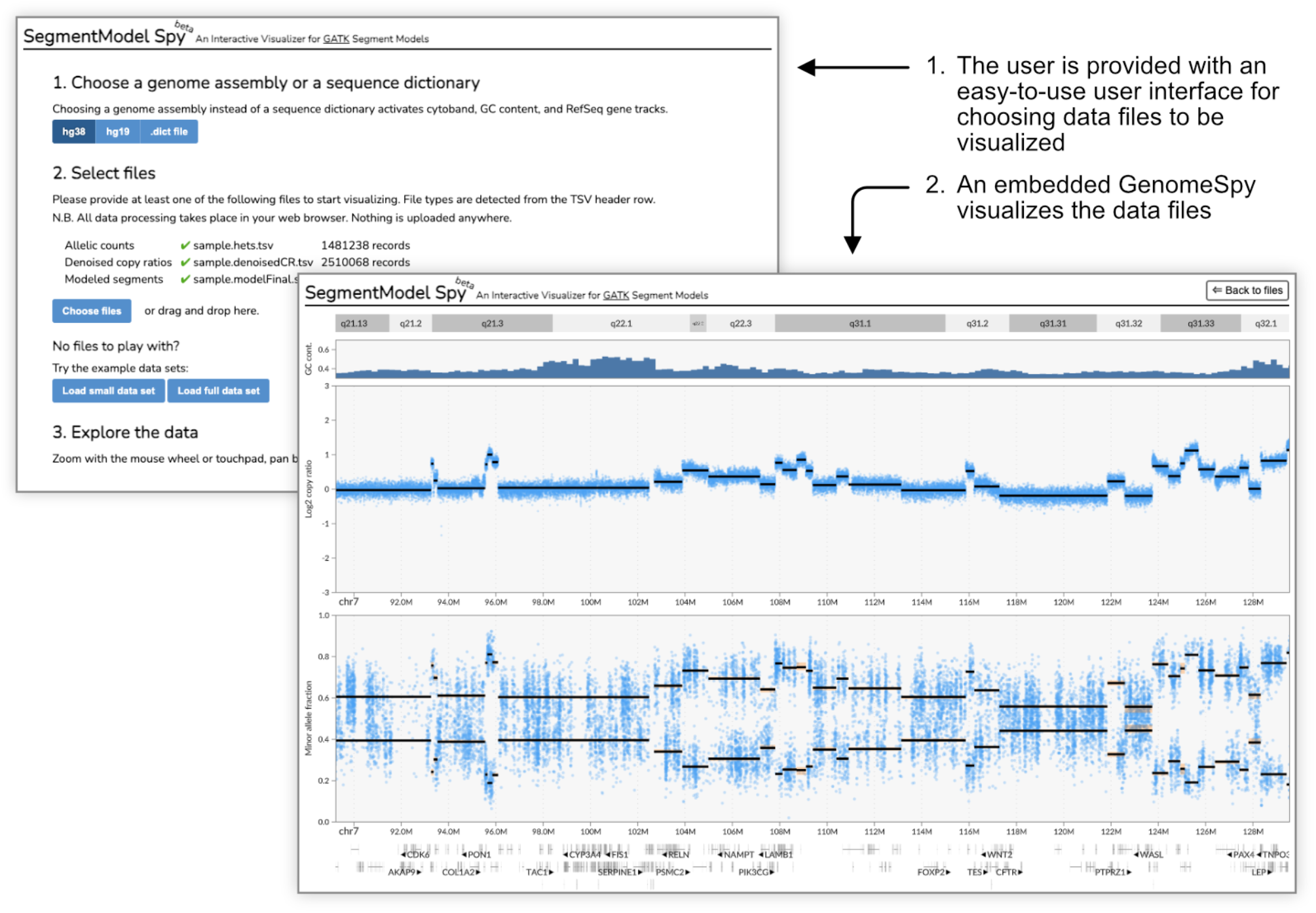
SegmentModel Spy demonstrates GenomeSpy’s utility as a visualization library in JavaScript web applications. It is a simple, end-user-oriented application for analyzing GATK’s copy-number segmentation results, allowing users to open data files effortlessly for swift navigation and inspection. The application generates a visualization specification and passes it with the parsed data files to the embedded GenomeSpy core library for visualization. Notably, all data processing occurs in the user’s web browser without the involvement of a remote server, enabling the analysis of sensitive data. SegmentModel Spy is available at https://genomespy.app/segmentmodel/.

The *app* is a general-purpose analytics application for large sample collections, built upon the core library (Figure 1C). It permits interactive analysis of genomic data and metadata, such as clinical variables. Using the grammar, users can adapt the app for different data types and analysis tasks. The app allows storing its state in the form of bookmarks or shareable links. The state comprises current scale domains, *i.e.*, shown genomic regions and the visibility of configurable visualization elements. The state also captures the filtering, grouping, and sorting actions performed on the samples, serving as provenance information that allows the recipient of a shared bookmark link to understand which steps led to a finding or insight [24, 25]. Finally, bookmarks also support optional Markdown-formatted notes, which allow communicating background information or implications related to the findings.

The *playground* web application (https://genomespy.app/playground/) integrates a code editor and a visualization, providing a convenient way to sketch new visualization designs. It is also the easiest method for new users to get started with GenomeSpy.

In addition to a specification, GenomeSpy visualizations need data, which can be provided as inline JavaScript objects in the specification or loaded from external files. CSV, TSV, and JSON files provide the highest flexibility. However, large data sets are better loaded lazily and only partially in response to user interactions, which is supported through compressed and indexed formats, such as BigBed, BigWig, FASTA, and GFF3 files. Additionally, the JavaScript API provides methods to dynamically update the data sets, enabling advanced use cases, such as integrations with and within other applications.

### Characterizing the genomic landscape of HGSC

We demonstrate the utility of the toolkit and highlight GenomeSpy App’s key features by showing how they enable the interpretation of WGS data from 753 samples of 215 patients belonging to the DECIDER clinical trial. Using GenomeSpy’s visualization grammar, we adapted the app for our data by specifying a visualization comprising segmented copy number alterations (CNA), loss of heterozygosity (LOH), somatic short variants (SSVs), and clinical data as shown in Figure 1C. We also specified several tracks exhibiting auxiliary information, such as ENCODE Blacklist [26], RefSeq Gene annotations [27], COSMIC Cancer Gene Census [28] and genes associated with platinum resistance [29]. Some of these tracks are hidden by default and can be activated from the toolbar. The visualization is available for exploration at https://csbi.ltdk.helsinki.fi/p/genomespy-preprint/.

#### Rapid transitions between the bird’s eye view and a closeup facilitates exploration

To streamline the exploration of large sample collections, we developed an interaction that rapidly transits the visualization from the bird’s eye view, which fits the whole collection into the available vertical space, to a close-up view, where the samples under the mouse cursor are shown in a larger size (Supplementary Video). This interaction allows for pinpointing interesting outliers among hundreds of samples and rapidly revealing them in sufficient detail for visual analysis, streamlining the exploration process. GenomeSpy’s GPU-accelerated rendering is pivotal in this feature, as it guarantees smooth transition between the views.

We used the bird’s eye view to gain an overview of the cohort. While recurrent *TP53* mutations and LOH on chromosome 17 (chr17) are known genomic aberrations in HGSC and contribute to tumor evolution [18, 30, 31], the concurrent display of both copy-number values and LOH revealed a striking pattern in the bird’s eye view: regardless of copy-number gains and losses in chr17, all but five patients presented a complete LOH in the whole chromosome (Figure 4). The whole-chromosome LOH suggests an early mitotic nondisjunction affecting the entire chromosome, with subsequent alterations, such as 17q amplifications, arising at a later stage.

**Figure 4:**
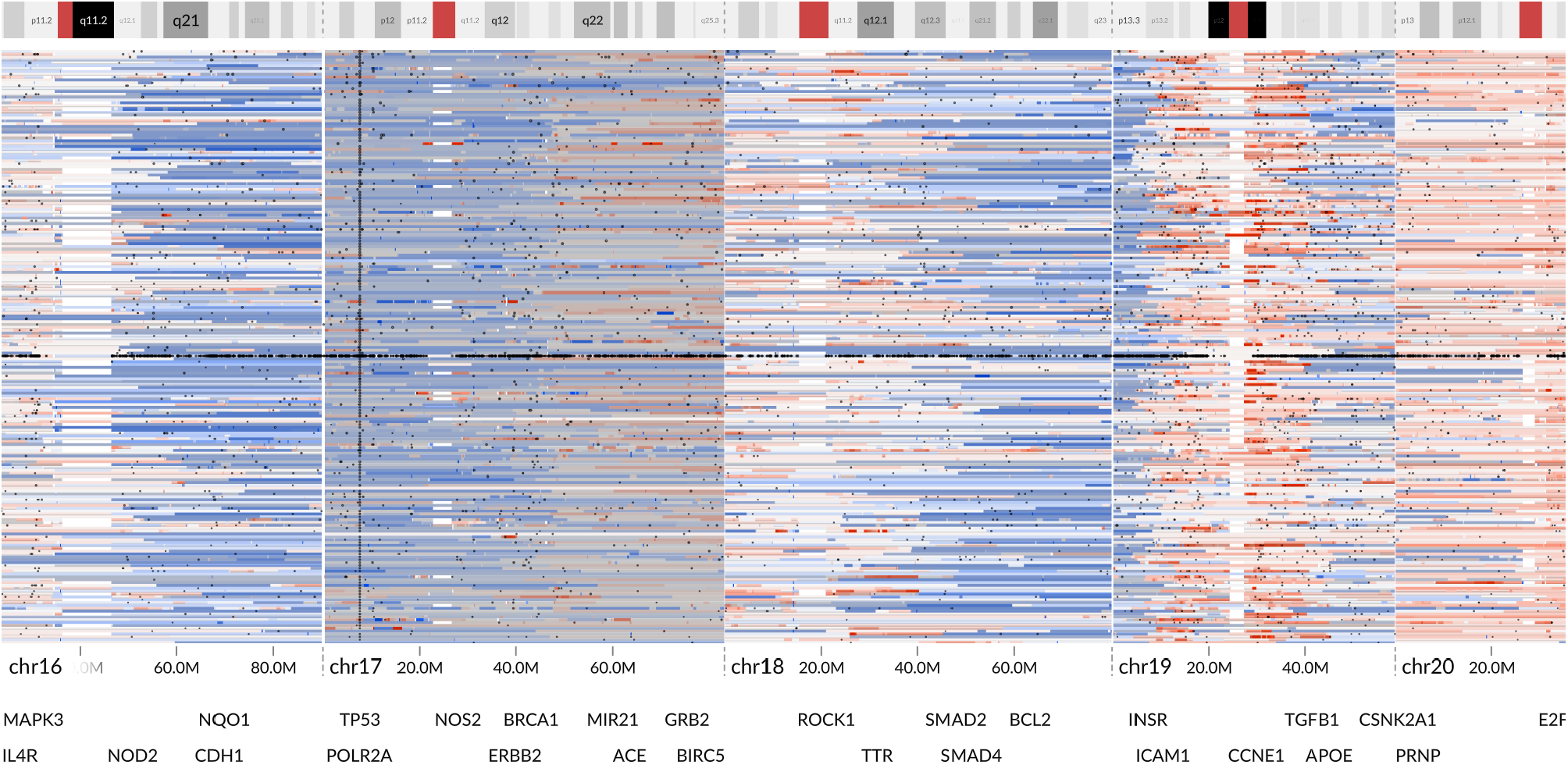
A bird’s eye view of all patients reveals a column of *TP53* mutations (dark dots) together with extensive LOH (gray overlay) on chr17. Only the sample with the highest purity (at least 15%) is included from each patient. One of the samples presents a very high number of SSVs and retained chr17 heterozygosity. The remaining four samples without full-chromosome LOH were from non-HGSC tumors. Link: https://csbi.ltdk.helsinki.fi/p/genomespy-preprint/#bookmark:TP53-and-LOH-in-chr17

We then looked more closely at the outliers that had retained chr17 heterozygosity by opening the close-up view (Supplementary video). Three of these outliers lacked a *TP53* mutation, which is atypical in HGSC. Thus, a gynecological pathologist re-evaluated these cases, and the diagnoses of the patients EOC466 and EOC545 were changed to low-grade serous carcinoma (LGSC) and EOC571 to endometrioid carcinoma. One of the outliers had lost heterozygosity only on 17p and was subsequently found to present endometroid carcinoma. The only HGSC tumor without chr17 LOH (patient EOC1106) stood out with a massive number of somatic mutations, indicating a possible mismatch-repair deficiency, which is a hallmark of Lynch syndrome. As Lynch syndrome results from germline mutations in DNA mismatch repair genes, we examined them and found a germline mutation in *MSH6*, which accounts for 10-20% of Lynch syndromes in colorectal cancer [32]. Since Lynch syndrome is dominantly inherited, these results were reported to a clinical geneticist to be discussed with the patient’s family.

#### Incremental, reversible actions enable rapid manipulation of the sample collection

Data exploration often involves the removal of irrelevant data items or organizing the data to uncover patterns. To facilitate this process, we developed a direct manipulation interface [33] that allows for incremental actions on abstract attributes such as clinical metadata or measurements at genomic loci. These actions can be accessed through a context menu (Figure 5), permitting the user to easily perform common tasks such as retaining samples belonging to a particular categorical class or stratifying samples based on a quantitative value at a specific genomic coordinate. Additionally, the actions are reversible, allowing for backtracking and further exploration of related questions. The actions also form a provenance record of the steps taken in the data exploration process, ensuring transparency and reproducibility.

**Figure 5:**
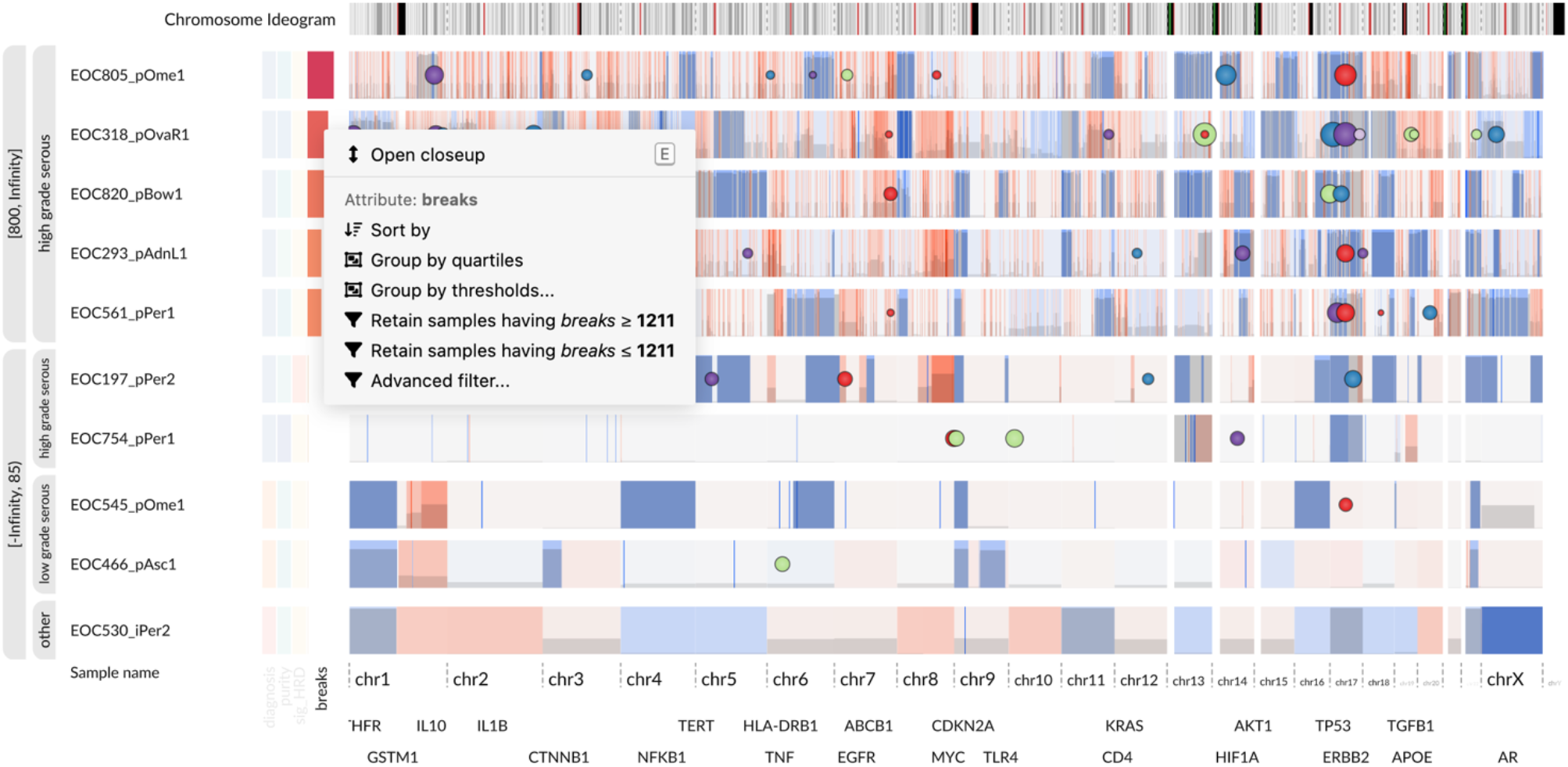
Top and bottom five samples by the number of copy-number breakpoints. Only the sample with the highest number of breakpoints was chosen from each patient. A nested, second-level grouping emphasizes the diagnosis attribute. The upper group exhibits a striking pattern of short amplifications associated with *CDK12* inactivation. The bottom group contains three samples from non-HGSC tumors and a peculiar HGSC tumor sample (EOC754_pPer1) with very few CNAs. The view was constructed using the incremental actions available through the attribute context menu (shown in the screenshot). Link: https://csbi.ltdk.helsinki.fi/p/genomespy-preprint/#bookmark:High-and-low-number-of-breakpoints

HGSC is characterized by extensive copy-number aberrations [18]. However, we observed considerable variation in the number of copy-number breakpoints between the patients. To better understand this variation, we applied a series of incremental actions to shape and stratify our sample set. First, we selected samples having purity at least 15%. We then sorted the samples into descending order by the number of breakpoints and retained the first, representative sample from each patient, which corresponded to the most fragmented one. Finally, we split the samples into groups based on the number of breakpoints and analyzed the patients with the most and least fragmented tumor genomes (Figure 5).

The five most highly fragmented samples showed a striking pattern of numerous focal amplifications evenly distributed throughout the genome. These amplifications ranged in size from ∼100kb to ∼10Mb. The zoomed-out whole-genome view also revealed deleterious (stop-gain or frameshift) *CDK12* SSVs (visible in chr17 in the figure) in four out of the five samples. The allele frequencies of the variants matched the tumor purity, suggesting homozygous mutations and thus, biallelic inactivation. Of note, four of these five samples presented copy-neutral LOH in the *CDK12* locus, suggesting subsequent amplification after the initial chr17 loss. Previous research has linked *CDK12* inactivation to a specific type of chromosomal instability characterized by tandem duplications with a bimodal size distribution, which is in line with our observation [34].

Interestingly, when visualizing all samples from these patients (Figure 6), the amplification pattern is nearly identical among the samples of each patient, implying subsequent stabilization of the genomes.

**Figure 6:**
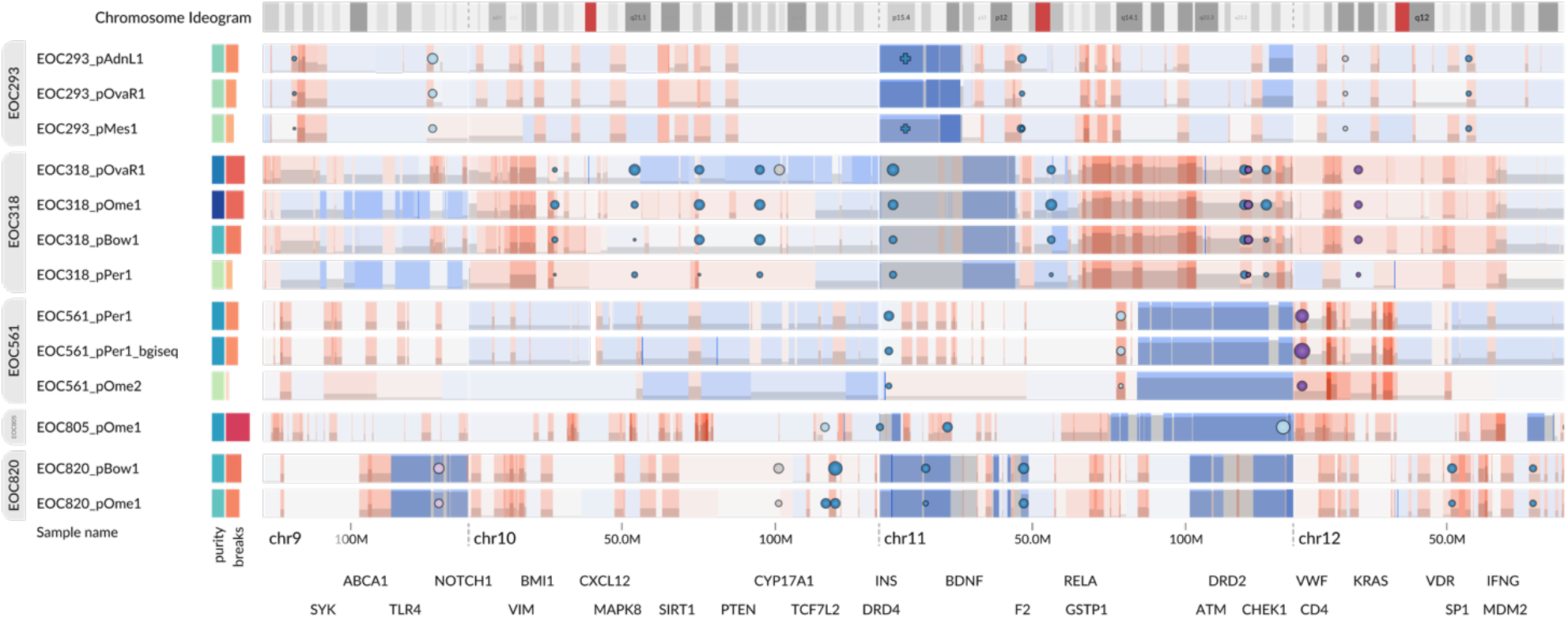
The amplified segments associated with the tandem-duplication phenotype and *CDK12* inactivation are largely identical within the samples of each patient, suggesting subsequent genome stabilization. Samples with a very low tumor purity, which are indicated by light green color in the metadata heatmap, suffer from low segmentation sensitivity and lack some of the segments that were detected in high-purity samples. Link: https://csbi.ltdk.helsinki.fi/p/genomespy-preprint/#bookmark:Top-5-fragmented-patients

Next, we focused on the five patients with the fewest breakpoints. Two of them (EOC466 and EOC545) were previously found to have LGSC based on the lack of a *TP53* mutation. Additionally, patient EOC530, who also lacked a *TP53* mutation but still exhibited chr17 LOH, had a non-serous neoplasm diagnosis. The two remaining patients had an HGSC diagnosis, but EOC754’s tumor presented a peculiar copy-number profile with aberrations only in three chromosomes. Although the mutated *TP53* and chr17 LOH in this tumor were consistent with the histological diagnosis of HGSC, the copy-number profile was surprising since it had even fewer arm-level aberrations than the two samples from LGSC patients.

We further analyzed the cohort for MAPK-pathway genes commonly altered in LGSC [35] and found *NRAS*:c.182A>G:p.Q61R in the samples from EOC530, EOC545, and EOC754, and another oncogenic aberration *BRAF*:c.1862A>G:p.N621S in samples from EOC466. Otherwise, oncogenic *NRAS* mutations were not detected in the entire cohort, and *BRAF* mutations were present in only two additional patients, EOC182 and EOC438, with characteristically simple copy number profiles. Generally, *NRAS* mutations are rarely seen in HGSC carcinomas but more commonly in borderline or low-grade serous neoplasms [36], as seen in patient EOC545.

As patient EOC754 exhibited an *NRAS* mutation and an atypical copy-number profile resembling the low-grade serous carcinomas of patients EOC545 and EOC466, a gynecological pathologist performed a retrospective histological review of her archival tumor samples. The tumor had a serous phenotype, but in terms of histological architecture, cytological atypia, and mitotic rate, the tumor, especially in ovarian samples, showed areas with unequivocally low-grade morphology in addition to areas with more pronounced pleiomorphism and mitotic activity. Yet, all four samples with sequencing data from this patient showed LOH on the whole chromosome 17 and a clonal *TP53* mutation in addition to *NRAS*. Cases with such genomic and morphological features from both high and low-grade serous carcinomas have previously been reported as rare variants of serous ovarian neoplasms [20, 37]. A further study on the potential origin and genomic and histological evolution of this and the two *BRAF*-mutated HGSC cases discovered through exploration in GenomeSpy is ongoing.

#### Score-based semantic zoom emphasizes important data items and mitigates overplotting

While somatic mutations are one of the driving forces behind tumorigenesis, most of the detected SSVs are passengers without contribution to disease. However, they clutter the view, making prompt identification of the pathogenic driver SSVs challenging. On the other hand, displaying all SSVs at once may be advantageous when an analyst studies a small genomic region that may accommodate SSVs with still uncertain pathogenicity. To address these conflicting needs, we developed *score-based semantic zoom*, a technique that couples a filter on an arbitrarily distributed quantitative attribute (*i.e.*, a score) with the zoom level (Supplementary Note, Supplementary Video). In the zoomed-out view, only the most important, *i.e.,* the highest scored data points, are shown, allowing the user to locate potentially important data items for a closer examination. As the user zooms in, items with lower scores become visible automatically, without the need to adjust separate filter settings. This behavior resembles online map applications where only the largest and most well-known place names are initially visible, with more names appearing gradually as the map is zoomed in. This technique also helps to avoid overplotting by controlling the number of concurrently visible data items.

To facilitate analysis and control overplotting, we applied the semantic zoom technique to all SSVs in the data set. For scoring, we used the Combined Annotation-Dependent Depletion (CADD) score [38], a single measure that integrates a diverse set of annotations. Thus, only the most likely pathogenic variants are shown at each zoom level. For instance, the recurrent *TP53* mutations and the *CDK12* mutations linked to chromosomal instability are visible already in the fully zoomed-out view (Figure 5), while the lower-scored variants remain out of sight until the user zooms in closer. This feature allowed us to instantly discover the pathogenic *CDK12* SSVs in the highly fragmented samples.

#### Data summarization allows easier comparison of stratified data

Although a CNA heatmap presents all details in data, a summary, such as the GISTIC G score [39], enables an easier perception of potential cancer driver regions and facilitates comparison of groups. Accordingly, we used GenomeSpy’s visualization grammar to specify a summary track that computes G scores over the segmented copy-number data. The summary incorporates a pipeline of transformations that inputs the copy-number values from the currently visible samples and computes a weighted coverage separately for amplifications and deletions (see Methods). A summary of the highest purity samples from all HGSC patients revealed a typical HGSC CNA landscape with prominent peaks around common HGSC driver genes [18], such as *MECOM*, *MYC*, *KRAS*, and *CCNE1* (Figure 7).

**Figure 7:**
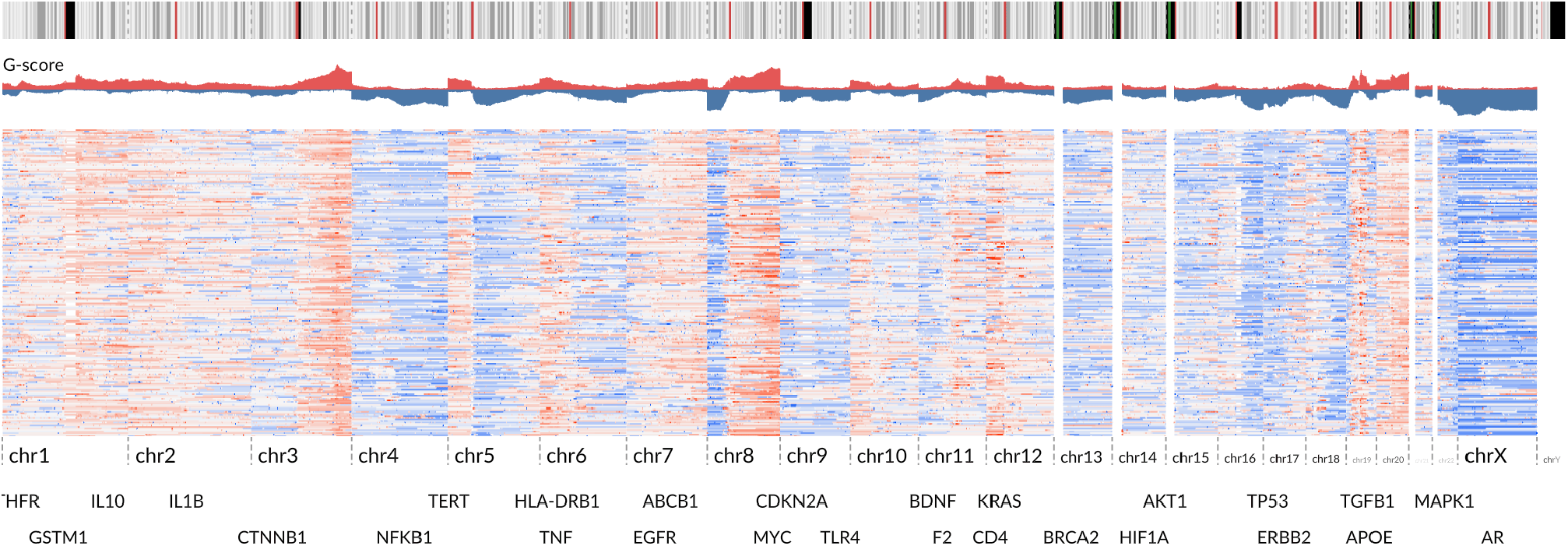
Using G score to summarize the copy-number landscape of the cohort. It is shown as an area chart above the copy-number heatmap. We used building blocks such as sample summarization, various transformations, and view compositions in the visualization specification to calculate and display the G score. Link: https://csbi.ltdk.helsinki.fi/p/genomespy-preprint/#bookmark:Copy-number-landscape

Next, we asked whether the recurrent amplification and deletion peaks in HGSC could be explained by clinical attributes or correlation of potential driver regions. Because the G-score summary track reflects the currently visible samples, and is computed separately for each group, we could easily analyze stratified data by visually comparing the G scores. However, stratifications failed to reveal distinguishable differences with attributes other than tumor ploidy. When we stratified the tumors into two groups approximately representing whole genome duplicated (WGD) and non-WGD tumors, an evident focal amplification peak around *CCNE1* in chr19 was present only in the WGD group, as shown in Figure 8. Previous research has associated such *CCNE1* amplifications with polyploidy and poor clinical outcome [40].

**Figure 8:**
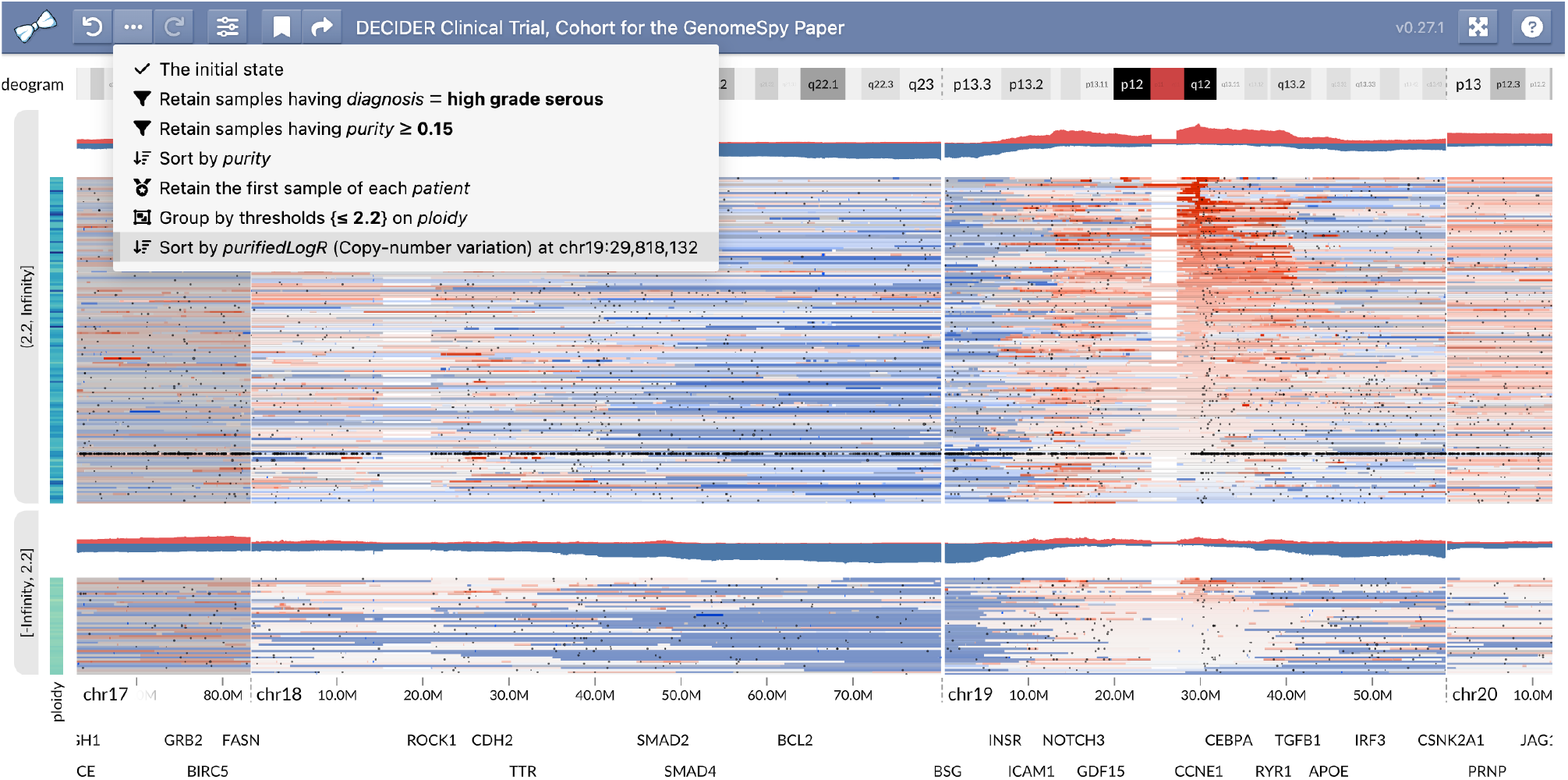
HGSC samples stratified by ploidy revealed higher *CCNE1* amplifications (shown in red) in the upper group that represents whole-genome-duplicated samples. Each group has a separately computed G-score summary to allow comparison. The opened drop-down menu reveals the provenance information comprising actions performed on the samples. Since most patients have samples from multiple tissues and time points, we kept only the highest-purity sample of each patient. Subsequently, we used the ploidy threshold of 2.2 to split the samples into two groups approximately representing non-WGD and WGD. Finally, we sorted the samples by the copy number of *CCNE1* to better illustrate the distributions of the copy-number log_2_(R) values in both groups. Link: https://csbi.ltdk.helsinki.fi/p/genomespy-preprint/#bookmark:WGD-and-CCNE1

### Data visualization helps in finding clinically actionable alterations

With the increased efforts to guide treatment decisions based on genomics findings, there is a need to rapidly visualize genomes to verify findings and detect aberrations that were not caught with automatic data analysis pipelines. For example, *BRCA1* is a tumor suppressor gene that contributes to DNA repair, and its mutation is an indication for targeted therapy with PARP inhibitors in HGSC [15].

As the PARP inhibitors are currently the only genomic-guided targeted therapy in HGSC, we used GenomeSpy to visually inspect the loci of *BRCA1* and other homologous recombination deficiency-related genes in our samples and identified a suspicious *BRCA1* region for the patient EOC763. A copy number pipeline, which employs GRIDSS [41] for joint structural-variation calling, confirmed a multi-exon in-frame deletion of *BRCA1* (chr17:43096222-43108182del, p.(K45_S198delinsN)) in all sequenced tumor samples from patient EOC763 (Figure 9). The deletion comprised exons 4-8, covering half of the RING domain. With supporting information from mutation signature analysis and the known consequences of similar medium-long deletions of *BRCA1* in ClinVar [42], we interpreted this *BRCA1* allele as pathogenic. Accordingly, the finding enabled the use of a PARP inhibitor to treat the patient in a recurrent setting. This example highlights the potential of visualization methods, such as GenomeSpy, in searching for genomically-based treatments for cancer patients.

**Figure 9:**
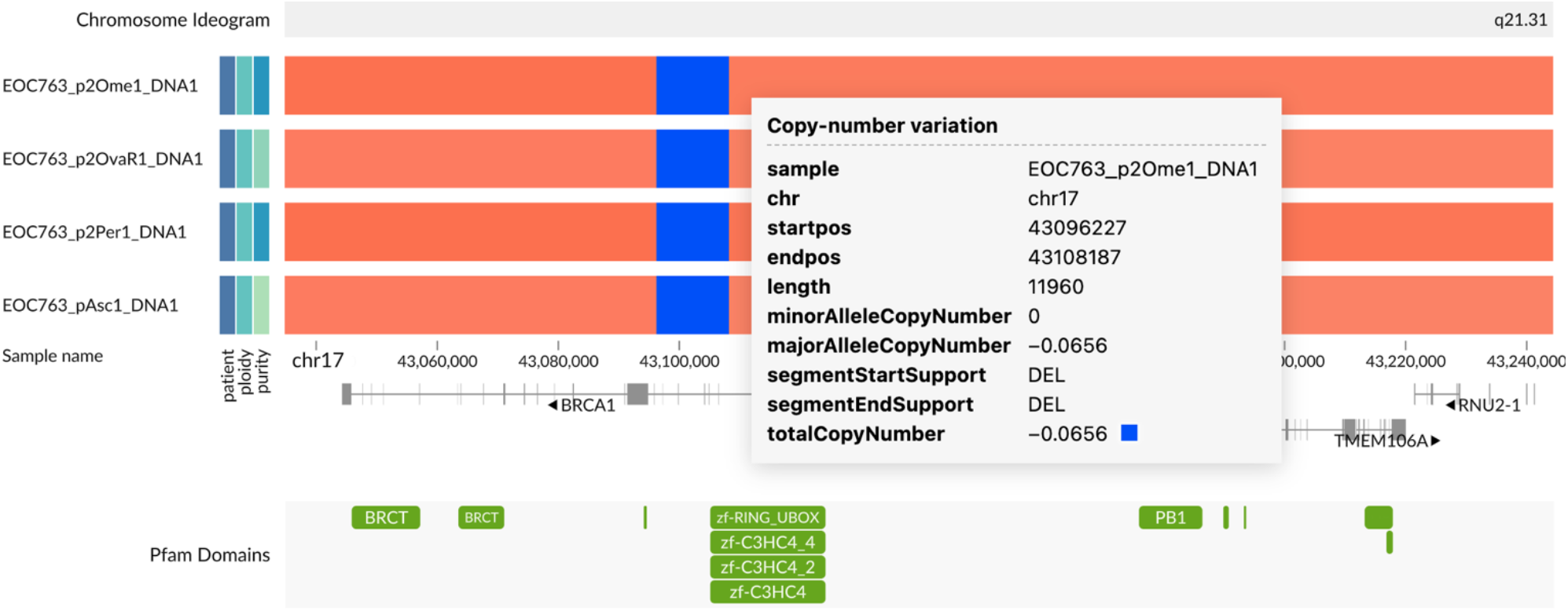
Results from an experimental copy-number pipeline revealed a homozygous *BRCA1* deletion in all tumor samples of patient EOC763. Because the pipeline could not directly output log_2_(R) and BAF values, we used the total copy number instead of log_2_(R) on the color channel of this visualization. Link: https://csbi.ltdk.helsinki.fi/p/genomespy-preprint/GRIDSS/

## Discussion

Visual exploration is a necessary step in oncogenomic data analysis and knowledge extraction [43]. To facilitate the exploration, we developed GenomeSpy, a visualization toolkit for genomic data. Two main objectives steered the process: designing a generic toolkit that enables effortless authoring of tailored visualizations for different use cases and implementing a fully customizable application to analyze large cancer sample collections. We achieved the genericity by implementing a grammar optimized for genomic data and demonstrated its expressivity, *i.e.*, its applicability to complex data, using the DECIDER cohort visualization. To support the swift analysis of sample collections, we applied the paradigm of fluid interaction [14], which manifested as several key features and influenced the overall architecture of the toolkit. For instance, we designed a GPU-accelerated rendering engine to allow rapidly updating graphics with an extensive number of data points. In addition to supporting continuous zooming and panning, it enabled the interaction that transits between the bird’s eye view and a closeup view, allowing quick examination of outliers in data sets. Importantly, its smooth animation helps the user stay focused without losing track of data not currently on the screen. Similarly, the score-based semantic zoom controls overplotting during navigation, allowing the user to focus on the most important data items at each zoom level. Finally, the direct-manipulation interface [33] based on incremental actions enables quick and versatile stratification and exploration with support for backtracking, bookmarks, and provenance information. All these features aim to expedite the exploration and thus foster insights.

GenomeSpy allowed us to characterize the genomic landscape of the DECIDER cohort, uncovering several interesting patterns. Among our findings, the extent and completeness of the chr17 LOH was one of the most surprising. Although such LOH has been previously found to occur in some ovarian carcinoma tumors [31], our data set shows that out of the 200 representative HGSC tumor samples having a purity of at least 15%, all but one, which had multiple *TP53* mutations, presented whole-chromosome LOH on chr17. While the whole-chromosome LOH allows a nascent tumor to expunge the remaining wild-type *TP53*, the same mechanism may also contribute to the biallelic inactivation or reduced dosage of other tumor-suppressor genes in the same chromosome, such as *CDK12*, *BRCA1*, *BRIP1*, and *NF1* [19, 44]. This hypothesis is supported by the pathogenic *CDK12* mutations associated with the tandem-duplicator phenotype. Homozygosity coupled with the copy-neutral LOH in these mutations indicates early occurrence, before or soon after the whole-chromosome loss. In addition to cohort characterization, GenomeSpy also enabled the discovery of exciting outliers, such as the tumor of patient EOC754 with traits from both HGSC and LGSC. Overall, the effective use of visual encodings and the high usability provided by fluid interaction have established GenomeSpy as an indispensable analysis tool among our geneticists, especially with copy number data, whose interpretation requires a view of the larger genomic context. Moreover, an example of using GenomeSpy to facilitate the identification of genomic-based personalized treatment is the discovery of an actionable *BRCA1* deletion, which was not detected with a commercial panel, most likely due to the small size of the deletion.

GenomeSpy visualizations allow end users, such as geneticists, clinicians, and bioinformaticians, to analyze data sets effortlessly. To make GenomeSpy accessible to a broader audience and to directly support use cases where established visualization designs already exist, we plan to furnish pre-defined visualization templates analogous to the common track types seen in genome browsers. Such templates will reduce the learning curve for new GenomeSpy users. Moreover, as the SegmentModel Spy example demonstrated (Figure 3), the toolkit can be used to build easy-to-use applications for specific analysis tasks. Expanding on this, we envision GenomeSpy as a foundation for a next-generation general-purpose genome browser that provides a comprehensive collection of data sets and pre-defined track types powered by extensive customizability and high-performance interactive graphics. Finally, although GenomeSpy’s grammar is already very expressive, our plans involve introducing additional building blocks. Examples of these include line and area marks, circular layouts, and parametrizable transformations, which all broaden the toolkit’s utility.

## Conclusions

In conclusion, we have demonstrated GenomeSpy’s flexibility and utility with the visualization of a cohort from the DECIDER clinical trial, and we envision the toolkit as a foundation for many future applications. The grammar-based approach allows its capabilities to be mixed and matched creatively, enabling tailored visualization in new research challenges. GenomeSpy is open-source software and is available together with documentation at https://genomespy.app/.

## Materials and Methods

### GenomeSpy Core

The core library is written in JavaScript. It uses the WebGL API and the TWGL library (https://twgljs.org/) for GPU-accelerated graphics. In addition, D3 [45] and Vega [46] libraries are used for CPU-side scale transformations, data loading, and expression handling. Genomic file formats, such as indexed FASTA, BigWig, BigBed, and GFF3 are loaded using GMOD JavaScript libraries [5]. The core library is available as an NPM package, which can be imported into web applications, web pages, and Observable notebooks. A more detailed description of the architecture and visualization grammar is available in the Supplementary Note and the GenomeSpy website (https://genomespy.app/).

### GenomeSpy App

The app builds upon the core library. It uses the Redux Toolkit (https://redux-toolkit.js.org/) for state management and provenance tracking and Lit (https://lit.dev/) for user-interface components. The application is available as an NPM package, which can be embedded on web pages together with a visualization specification and data.

### DECIDER Cohort

“Multi-layer Data to Improve Diagnosis, Predict Therapy Resistance and Suggest Targeted Therapies in HGSOC” (DECIDER; ClinicalTrials.gov identifier: NCT04846933) is a prospective, longitudinal, multiregion observational study that began recruitment in 2012. Herein, we included 215 patients treated at Turku University Hospital, Finland. The treatment was either primary debulking surgery (PDS), followed by a median of six cycles of platinum-taxane chemotherapy, or neoadjuvant chemotherapy (NACT), where primary laparoscopic operation with diagnostic tumor sampling was followed by three cycles of carboplatin and paclitaxel.

Altogether we included all 753 tumor samples that had been whole-genome-sequenced when the cohort was formed. The samples comprise tumor tissue (tubo-ovarian, intra-abdominal, and other metastatic sites such as lymph nodes) and ascites from several phases of the disease.

All patients participating in the study gave their informed consent. The study and the use of all clinical materials have been approved by the Ethics Committee of the Hospital District of Southwest Finland (ETMK) under decision number ETMK: 145/1801/2015.

### Whole-Genome Sequencing

Genomic DNA was extracted from tumor tissue or ascites cells and whole blood or buffy coats isolated from whole blood. After assessing DNA quality, the samples were whole-genome sequenced with either DNBSEQ (BGISEQ-500 or MGISEQ-2000, MGI Tech Co., Ltd., China), NovaSeq 6000 (Illumina, USA), or HiSeq X Ten (Illumina, USA) as 150bp paired end sequencing. Median coverage was ∼47x (range 23–158x). Raw read data were processed with Trimmomatic [47], FastQC (https://www.bioinformatics.babraham.ac.uk/projects/fastqc/) in the Anduril 2 workflow platform [48]. The reads were then aligned to the human genome GRCh38.d1.vd1 using BWA-MEM, followed by a duplicate removal with Picard Tools (http://broadinstitute.github.io/picard/) and base quality score [49] recalibration with the Genome Analysis Toolkit (GATK) [50].

### Mutation Calling

We called somatic mutations using GATK *Mutect2* [51] with joint calling [52]. A panel of normals generated from 181 DECIDER and 99 TCGA blood-derived normal samples was used. Mutations were annotated using ANNOVAR [53], ClinVar [42], and CADD estimates for deleteriousness [38]. Germline mutations were jointly called using GATK [52] from 217 DECIDER normal samples with allele-specific variant quality score recalibration. Variant quality score recalibration was allele specific. Mutational signatures were fitted using COSMIC v3.2 signatures [54, 55], adjusted for GRCh38 nucleotide frequencies.

### Copy-Number Calling and Estimation of Ploidy and Tumor Purity

We used GATK to perform the copy-number segmentation. The analysis pipeline follows the GATK best-practices documentation and builds upon the Anduril 2 platform.

To collect the minor allele counts (b-allele frequency, BAF), we used all filtered biallelic germline SNPs with heterozygous calls (VAF between 40% and 60%) from each patient. Both read and allelic count collection excluded regions listed in the ENCODE blacklist [26] and our internal DECIDER blacklist, which is available as a track in the DECIDER visualization. The DECIDER blacklist includes regions having abs(log_2_(R)) > 0.2 in at least three of the 114 normal samples used as input data. The 136 regions in the DECIDER blacklist represent poorly aligned regions and population-level copy-number variance. We used platform-specific (HiSeq, DNBSEQ, and NovaSeq) panels of normals built from the normal samples to denoise the read counts.

Since the result of the actual segmentation affects downstream analyses such as ploidy and purity estimation, we visually evaluated the effect of the various parameters of GATK’s *ModelSegments* tool. In practice, we ran the segmentation for select samples using 729 different combination of values for the parameters and studied their effect using the SegmentModel Spy tool (Figure 3, Supplementary Note). Finally, we chose parameters that resulted in the subjectively best breakpoint inference results. For instance, short segments should be included, but false breakpoints related to GC-wave artifacts need to be avoided. The final parameters were as follows: number-of-changepoints-penalty-factor: 1, kernel-variance-allele-fraction: 0, kernel-variance-copy-ratio: 0.2, kernel-scaling-allele-fraction: 0.1, smoothing-credible-interval-threshold-allele-fraction: 2, smoothing-credible-interval-threshold-copy-ratio: 10.

After the segmentation, we used a reimplemented ASCAT algorithm [56] to estimate purity, ploidy, and allele-specific copy numbers. The original ASCAT R package was not directly applicable because it fails to accept data segmented using external tools. Our implementation also uses the variant-allele frequency (VAF) of truncal pathogenic *TP53* mutation as additional evidence in selection of the optimal ploidy/purity solution. As nearly all patients have a homozygous *TP53* mutation in their cancer cells, we can use the VAF and the estimated total copy number (CN) of *TP53* to approximate the purity:

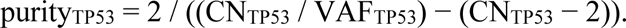

Patients having discordant ploidy estimates between their samples went through manual curation.

Since the contribution of non-aberrant cells on the log_2_(R) and BAF values encumber visualization and further analyses, we calculated “purified” values, *i.e.*, what the log_2_(R) would be in the absence of normal cells.

Purified R, based on discussion in https://github.com/lima1/PureCN/issues/40:

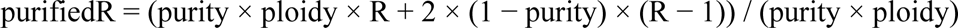

Purified BAF, derived from S2, S7, and S8 of (van Loo et al., 2010):

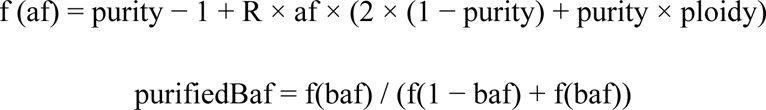

### Experimental Copy-Number Pipeline for *BRCA1/2* Analysis

We called structural variants in a callset of 139 DECIDER patients using GRIDSS [41] with joint calling and performed the somatic filtering using GRIPSS (https://github.com/hartwigmedical/hmftools/tree/master/gripss) with a panel of normals from Dutch population [57] and the ENCODE blacklist [26]. The BAF was calculated using AMBER (https://github.com/hartwigmedical/hmftools/tree/master/amber) with the heterozygous SNP loci from the mutation calling. Read depth was extracted using COBALT (https://github.com/hartwigmedical/hmftools/tree/master/cobalt), which also performed GC normalization. Finally, we employed PURPLE [57] to combine BAF, read depth ratios, and structural variants to estimate purity, ploidy, and the copy-number profile of the samples.

### Pathogenic *BRCA1/2* Mutations

We curated somatic and germline short variants in *BRCA1/2* genes. We considered a variant pathogenic, if it causes premature truncation in the canonical transcript or if it is annotated as pathogenic or likely pathogenic in the ClinVar [42] database. For patient homozygosity assessment, we compared allelic read counts against allele-specific copy numbers in the locus and purities in tumor samples with a minimum purity of 5%. A variant was considered homozygous, if it was the most likely explanation for the allelic read counts across a patient’s tumor samples.

### DECIDER Cohort Visualization

We used the GenomeSpy app for the DECIDER visualization. Annotation tracks such as RefSeq genes are specified in separate JSON files, allowing easy reuse. The main JSON file specifies the visualization of metadata, SSVs, CNV, BAF, and the copy-number summary. GenomeSpy inputs all genomic and metadata from tab-separated values (TSV) files.

Only SSVs with the CADD score of at least 10.0 or that were pathogenic according to ClinVar (Landrum et al., 2018) were included to reduce loading time and memory consumption. We used the purified log_2_(R) and BAF values for CNV and LOH, allowing more meaningful comparison, sorting, and grouping. To enable easier perception of aberrant BAF, we converted it into LOH using the formula:

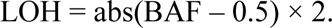

Here, zero indicates full heterozygosity, one indicates a total loss of heterozygosity.

The dynamically updating copy-number summary track replicates the G-score of GISTIC 1.0. Briefly, the dataflow processes amplifications and deletions separately. Only segments with abs(purifiedLogR) > 0.1 are included and abs(purifiedLogR) is clamped to 1.5. Finally, the dataflow computes a purifiedLogR-weighted coverage for the segments and divides it by the number of samples involved. Coverages of amplifications and deletions have separate layers in the visualization and are shown as red and blue, respectively.

The RefSeq gene annotation track uses a popularity-based prioritization for the gene symbols [58], a method originally introduced in HiGlass [59]. Thus, at each zoom level, the symbols are handled in priority order and shown if there is still room on the track.

## Abbreviations

BAF: B-allele frequency
CNV: Copy number variance
DECIDER: Multi-layer Data to Improve Diagnosis, Predict Therapy Resistance and Suggest Targeted Therapies in HGSOC
GPU: Graphics processing unit
JSON: JavaScript Object Notation
LOH: Loss of heterozygosity
PARP: ADP ribose polymerase
SSV: Somatic short variant
VAF: Variant allele frequency
WGS: Whole-genome sequencing

## Declarations

### Code Availability

Project name: GenomeSpy

Project home page: https://genomespy.app/

Operating systems: Not applicable

Programming languages: JavaScript and TypeScript

License: MIT

### Availability of Data and Materials

- The GenomeSpy toolkit: https://github.com/genome-spy/genome-spy (https://doi.org/10.5281/zenodo.7852282)
- SegmentModel Spy: https://github.com/genome-spy/segment-model-spy
- The DECIDER HGSC visualization is available for exploration at: https://csbi.ltdk.helsinki.fi/p/genomespy-preprint/
- The visualization specification and processed data are available at: https://csbi.ltdk.helsinki.fi/p/genomespy-preprint/spec.json (will be submitted to a repository by the time of publishing)
- All sequencing data will be available at the European Genome-phenome Archive (EGA) under accession number EGAS00001006775.
- Any additional information required to reanalyze the data reported in this paper is available from the lead contact upon request.

### Author Contributions

KL, JO, and RL conceptualized the project. KL designed and implemented the software and all visualizations. SHa supervised the project. KL, JO, YL, TM, GMi, GMa, and AL analyzed sequencing data. AV performed histopathological analysis. KH, SHi, and JH acquired samples. KL wrote the manuscript with contributions from YL, TM, GMi, AV, and SHa and feedback from all authors.

## Acknowledgements

The authors acknowledge CSC-IT Center for Science, Finland, for computational resources. ChatGPT and Grammarly were used to improve the grammar, vocabulary, and the flow of the text.

## Ethics Declarations

### Ethics approval and consent to participate

The study and the use of all clinical materials have been approved by the Ethics Committee of the Hospital District of Southwest Finland (ETMK) under decision number ETMK: 145/1801/2015.

### Consent for publication

Not applicable.

