## Supplementary Note for "Deciphering Cancer Genomes with GenomeSpy: A Grammar-Based Visualization Toolkit"

### GenomeSpy: Supplementary Notes

### 1. Comparison to Vega-Lite

The visualization grammar of GenomeSpy is heavily inspired by Vega-Lite [1]. However, Vega-Lite lacks support for genomic data, and the rendering performance of Vega [2], which serves as an "engine" for Vega-Lite, is insufficient for large genomic datasets. Still, its overall design that combines graphical marks, scales, transforms, view composition, and data input into a concise grammar is a suitable starting point for a more domain-specific grammar. We replicated the most useful parts of Vega-Lite and extended it with features that are necessary when working with genomic data. GenomeSpy provides partial compatibility with Vega-Lite's view specifications because we have intentionally avoided reinventing or challenging design decisions made in Vega-Lite (Supplementary Figure 1). Thus, GenomeSpy should be very familiar to those with experience working with Vega-Lite. Next, we describe the main similarities and differences between the two. Full documentation is available at https://genomespy.app/.

#### Handling Genomic Coordinates

Genomic coordinates are commonly presented as tuples comprising a discrete chromosome and an integer position within the chromosome. To allow for straightforward navigation around the whole genome, the discrete chromosomes could be concatenated into a single linearized coordinate system (Supplementary Figure 2a).

GenomeSpy provides a specific `linearizeGenomicCoordinate` transform that performs the linearization. However, because handling genomic coordinates is such a common task in genome visualization, GenomeSpy also allows for specifying the chromosome and position fields directly in the `encoding` block (Supplementary Figure 2b).

Behind the scenes, GenomeSpy inserts a `linearizeGenomicCoordinate` transform into the data flow and replaces the `chrom` and `pos` properties with a reference to the newly computed linearized field.

Linearization needs the chromosome sizes and their order, which can be provided as a `chrom.sizes` file downloadable from the UCSC website. Alternatively, they can be inlined directly into the visualization specification. All the necessary files for `hg38`, `hg19`, `hg18`, `mm10`, `mm9`, and `dm6` assemblies

```
{
  "width": 250,
  "height": 200,
  "data": {
    "values": [
      { "a": "A", "b": 28 }, { "a": "B", "b": 55 },
      { "a": "C", "b": 43 }, { "a": "D", "b": 91 },
      { "a": "E", "b": 81 }, { "a": "F", "b": 53 }
    ]
  },
  "encoding": {
    "x": {
      "field": "a",
      "type": "nominal",
      "scale": { "padding": 0.1 },
      "axis": { "labelAngle": 0 }
    },
    "y": { "field": "b", "type": "quantitative" }
  },
  "layer": [
    { "mark": "rect" },
    {
      "mark": { "type": "text", "dy": -9 },
      "encoding": {
        "text": { "field": "b" }
      }
    }
  ]
}
```

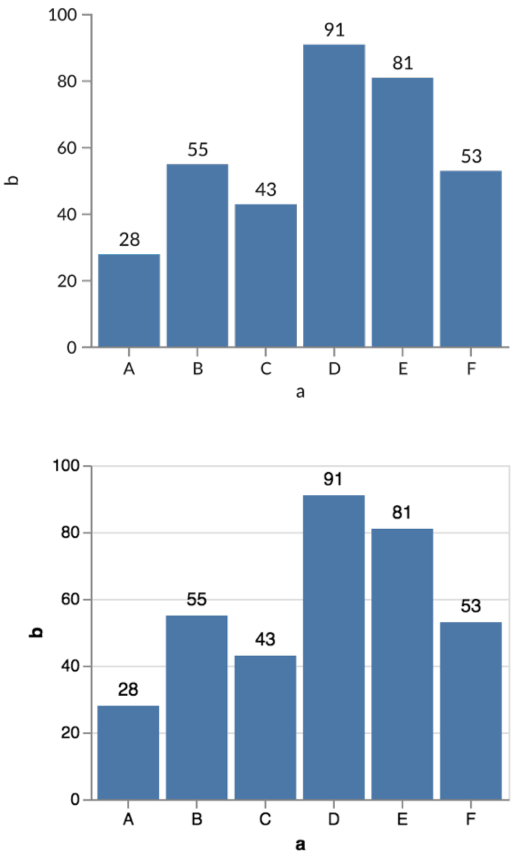

Supplementary Figure 1: An example of a visualization specification with two layers rendered in GenomeSpy (top) and Vega-Lite (bottom). GenomeSpy adapts the design of Vega-Lite's grammar, providing partial compatibility. However, the implementation is independent and makes extensive use of GPU in scale transformations and rendering. In this example, the data set is embedded into the specification and comprises objects with two fields: a and b. The *encoding* block specifies how the data fields are mapped to different visual channels. In this case, the a field is declared as nominal data and mapped onto the x axis, and the y field is quantitative and mapped onto the y axis. The *layer* block specifies two superimposed graphical marks: *rect* forms the bars on the chart and *text* shows the exact data values above the bars.

are readily available and loaded automatically from the https://genomespy.app web server by specifying the name of the genome assembly in the visualization specification.

Documentation for genomic coordinates: https://genomespy.app/docs/genomic-data/genomic-coordinates/

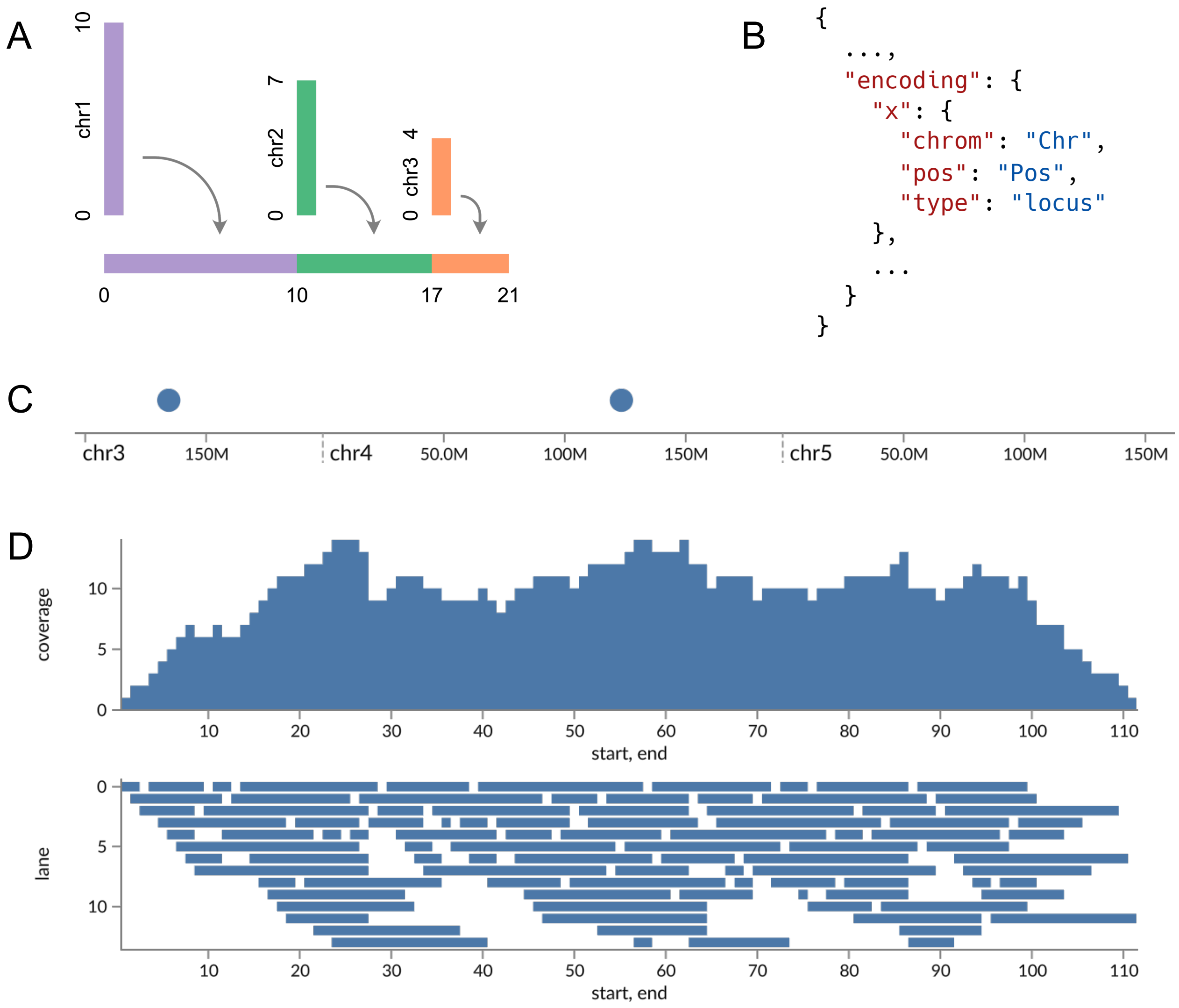

Supplementary Figure 2: Handling genomic data in GenomeSpy using transformations and scales. (A) In an abstract sense, a transformation inputs a list of data items and outputs a list of new items that may be filtered, modified, or generated from the original items. GenomeSpy provides a *linearizeGenomicCoordinate* transformation that maps the discrete chromosomes onto a single linear coordinate space. (B) By using the *locus* data type and specifying the *chrom* and *pos* fields, an implicit linearization transformation is added to the data flow, allowing easy handling of genomic coordinates. (C) The *locus* scale maps the linearized genomic coordinates to the viewport and provides chromosome-aware axis ticks. (D) GenomeSpy provides several data transformations that enable visualization techniques commonly used with genomic data. For example, when working with overlapping segments, the *coverage* transformation (upper plot) generates a list of new segments with continuous coverage values and the *pileup* transformation (lower plot) assigns each segment a free lane. Both plots use the *rect* mark to visualize the transformed data items.

#### Scales

GenomeSpy implements most Vega-Lite's scales, including `linear`, `pow`, `sqrt`, `symlog`, `log`, `ordinal`, `band`, `point`, and `threshold`. However, none are particularly suitable for genomic or amino acid coordinates, which are discrete and indexed using integers. Therefore, we extended the grammar with two new scales: `index` and `locus`.

The `index` scale is designed for zero-based, half-open coordinates used in the UCSC-family file formats such as `BED`. The scale renders similarly to the discrete `band` scale but behaves like the `linear` scale; the scale domain comprises an arbitrary interval of indices, and it can be freely zoomed and panned. Index scale is suitable for data that represents, for example, a single chromosome or a specific nucleotide sequence.

The `locus` scale is identical to `index` scale but provides a chromosome-aware axis. When working with linearized coordinates, `locus` scale displays chromosome labels and intra-chromosomal positions on the axis (Supplementary Figure 2c).

Documentation for scales: https://genomespy.app/docs/grammar/scale/

#### Transformations

GenomeSpy supports many of the transformations of Vega-Lite, including but not limited to `filter`, `formula`, and `stack`. We also implemented several new transformations that support common genome visualization tasks. For example, `coverage` transformation computes coverage for overlapping genomic ranges, and `pileup` transformation computes a piled-up layout for such ranges (Supplementary Figure 2d).

Documentation for transformations: https://genomespy.app/docs/grammar/transform/

#### Marks

GenomeSpy provides `point`, `rect`, `rule`, and `text` marks, which are largely compatible with their Vega-Lite counterparts. We extended the `text` mark with support for secondary positional coordinates, which enables useful behaviors for the rendered text. For instance, the text can be shrunken so that it always fits the interval, or it can be geometrically translated so that it remains inside the viewport even if either one of the endpoints resides outside the viewport. These behaviors are particularly useful when annotating genomic regions, such as cytobands. The secondary positional channel in text mark also enables sequence-logo visualizations (Figure 1b).

We also implemented a `link` mark, which displays arcs between two endpoints (Figure 1b), enabling structural-variant visualizations.

Documentation for marks: https://genomespy.app/docs/grammar/mark/

#### View Composition

The GenomeSpy core library provides `concat` (vertical, horizontal, and wrapping concatenation) and `layer` compositions, which enable diverse visualization layouts. For example, vertical concatenation with shared scales on the `x` channel allows the creation of track-based layouts that mimic genome browsers.

The GenomeSpy app extends the available compositions with sample-based faceting that creates an own sub track for each sample. The app also provides various interactions that allow the end user to filter, sort and group the samples. Data can be further aggregated into summary tracks that are computed separately for each group.

Documentation for concat and layer: https://genomespy.app/docs/grammar/composition/
Documentation for sample-based faceting: https://genomespy.app/docs/sample-collections/visualizing/

#### Data Input

Comma/tab-separated values (CSV, TSV) and JSON are supported out of the box because GenomeSpy uses the `vega-loader` package from Vega for data input. We also implemented a FASTA loader, which can be used, for example, in multiple-sequence alignment visualizations.

Documentation for data input: https://genomespy.app/docs/grammar/data/

#### Interactivity

Vega-Lite's interactivity is based on the parameter/selection concept, which builds upon the reactive streams and signals of Vega. Albeit flexible, it is an unnecessarily complex architecture for basic interactions such as zooming and panning. In GenomeSpy, scales on positional channels can be made zoomable using the `zoom` property of the scale properties.

#### Omitted Vega-Lite features

Because we developed GenomeSpy mainly for use cases involving genomic data, some Vega-Lite functionality, such as support for temporal and geographical data, have been entirely omitted.

#### 2. GPU-accelerated architecture

The GenomeSpy core library uses the WebGL 2.0 API to access the GPU from JavaScript. WebGL is a low-level API that allows transferring data between the main and GPU memory, drawing triangles, and performing vertex and pixel manipulation using shader programs written in the GLSL language (https://webglfundamentals.org/webgl/lessons/webgl-fundamentals.html).

GPU access in GenomeSpy is handled by the mark classes (`rect`, `point`, etc.). They build mark-specific vertex buffers from the raw data and dynamically generate shader code that maps the raw data values to visual channels using scale functions. In practice, marks serve as a high-level abstraction on the low-level WebGL API. For example, we built complex visualization elements such as axes from the marks using the visualization grammar. This way, we avoided introducing any WebGL and shader code that would be specific to the axes.

The actual rendering comprises two phases. The layout phase traverses the view hierarchy, calculates view coordinates, and builds an optimized rendering batch. It also reorders the batched operations to minimize WebGL state changes between draw calls, significantly improving performance with large facet quantities. The actual rendering phase executes the batch, which mainly performs WebGL API calls that change shader programs, set uniform values such as scale domains, and perform draw calls. When the user interacts with the visualization by zooming or panning, scale domains are adjusted accordingly, and only the rendering phase is re-executed. Particularly, vertex buffers are updated between frames only if new data is fed to the visualization. The highly optimized rendering phase ensures minimal CPU utilization, a high frame rate, and smoothly animated interactions.

The rendering architecture of GenomeSpy has similarities to Stardust [3] and P4 [4], both of which use the GPU for scale transformations. In contrast, Vega uses the central processing unit (CPU) to execute a reactive dataflow graph, which produces a scene graph comprising all mark instances. The scene graph is subsequently rendered into an HTML canvas or SVG graphics. Although a WebGL renderer exists for Vega (https://github.com/vega/vega-webgl-renderer), it fails to provide a significant performance improvement over the canvas renderer: when the scene graph is altered – for example, when the user zooms in a scatter plot – scene graph items must be updated to vertex buffers and reuploaded into GPU memory. In conclusion, although GenomeSpy's visualization grammar resembles

Vega-Lite, the rendering architecture is highly dissimilar, optimized for navigating large genomic datasets.

#### 3. Embedding in web applications: SegmentModel Spy

This section demonstrates the usage of GenomeSpy as a visualization library in a special-purpose web application. Certain cancers exhibit high amounts of copy-number aberration [5], which results in altered gene expression through loss or amplification of genomic regions. An essential part of copy-number analysis is segmentation, i.e., the extraction of continuous regions from noisy raw data, consisting of read and allelic counts in small, commonly one kilobase windows.

The Genome Analysis Toolkit (GATK) [6] is one of the available methods for copy-number segmentation. It provides a built-in *PlotModeledSegments* (https://gatk.broadinstitute.org/hc/en-us/articles/360042477992-PlotModeledSegments) tool, which produces static plots of the raw data overlaid by the estimated segments. However, because the visualization is static and comprises the whole genome, a comprehensive assessment of the estimated segmentation breakpoints is difficult to achieve. To combat this problem, we developed *SegmentModel Spy* (https://github.com/genome-spy/segment-model-spy), an interactive visualization tool, using the core GenomeSpy library. The tool is an easy-to-use web application that replicates the output of the *PlotModeledSegments* tool but provides interactivity with continuous zooming and panning, allowing for an accurate evaluation of results and the adjustment of the segmentation parameters.

The user launches the visualization by choosing an optional genome assembly and dragging and dropping any combination of GATK data files (such as read/allelic counts or segmentation) to the application window. All data processing takes place in the user's web browser, allowing for the viewing of sensitive data. Once the application has parsed the data files, the visualization appears (Figure 3). The user can zoom in/out with the mouse wheel and pan the view by dragging. Alternatively, the user can use touchpad gestures. Because the GPU-accelerated rendering ensures smooth and rapid interactions, the user can swiftly scan through the segmentation results and quickly focus on the critical details, such as missed segments.

If the user chooses one of the supported genome assemblies, the view is augmented with annotations such as cytobands and RefSeq gene annotations, which provide context for the copy-number segments. In addition, as the read counts often contain GC-wave artifacts that interfere with the segmentation [7], the visualization includes a track that displays GC content in 250 kilobase windows. Thus, the user can consider whether a segmentation breakpoint results from an artifact or an actual signal in the data.

Technically, SegmentModel Spy uses the GenomeSpy core library for visualization. SegmentModel Spy handles the file selection interface, parses the data files, and feeds the data along with a visualization specification to the GenomeSpy core for displaying. With four million data points on a MacBook Pro (13-inch, 2018) and a Radeon Pro 580 eGPU, the visualization can maintain a constant 60 frames-per-second performance at all zoom levels.

### 4. Exploring sample collections with the GenomeSpy App

The *app* package provides a framework for analyzing large sample collections. It provides no ready-made visualizations – instead, the user uses the visualization grammar to specify a visualization for a data set. The app extends the grammar by providing faceting for multiple samples, which are shown as separate tracks in the visualization. Moreover, users can manipulate the sample collection with various actions that the app provides, enabling complex analyses. A comprehensive description of the features is available in the documentation.

#### Rapid transitions between the bird's eye view and a closeup

To streamline the exploration of large sample collections, we developed a novel interaction that transits the visualization from the bird's eye view, which fits the whole collection into the available vertical space, to a close-up view, where the samples under the mouse cursor are shown in a larger size (Supplementary Video). The interaction allows for pinpointing interesting outliers among hundreds of samples and rapidly revealing them in sufficient detail for visual analysis. The usability of the interaction depends on the smooth transition animation between the two states: the animation allows the user to mentally retain the context of the off-screen samples, which would be difficult with an abrupt or choppy transition.

Documentation for the app: https://genomespy.app/docs/sample-collections/

#### 5. Comparison to Gosling

Gosling [8] is another grammar-based visualization toolkit for genomic data, designed for interactive visualizations running in a web browser. Table 1 summarizes the main differences between Gosling and GenomeSpy.

Table 1: Comparison of Gosling and GenomeSpy

| Gosling | GenomeSpy |
|---|---|
| • More domain-specific<br>• Grammar more divergent from Vega-Lite<br>• Uses pixi.js (https://pixijs.com/) library for graphics abstraction<br>• Supports circular layouts<br>• Supports the brushing interaction<br>• Embeddable in JavaScript and Python notebooks<br>• Written in TypeScript | • More generic<br>• Grammar more similar to Vega-Lite (partial compatibility)<br>• Implements graphics abstraction using custom GPU shader programs optimized for the grammar<br>• Superior rendering performance<br>• Built-in support for the exploration and analysis of large sample collections<br>• Embeddable in JavaScript notebooks<br>• Written in JavaScript with JSDoc type annotations |

#### Performance comparison

This subsection presents a brief performance comparison of GenomeSpy and Gosling, highlighting the difference in CPU utilization stemming from different architectures. Supplementary Figure 3 and Supplementary Figure 4 show Chrome Developer Tool performance profiles for typical, non-trivial GenomeSpy and Gosling visualizations, run on a 2021 MacBook Pro (M1 CPU) and Chrome 116. Although the profiles represent entirely different visualizations, profiling other publicly available GenomeSpy and Gosling visualizations produces similar results. In brief, GenomeSpy makes more effective use of GPU, significantly reducing the CPU utilization when navigating data sets, and thus, ensures smooth animation during navigation and other interactions.

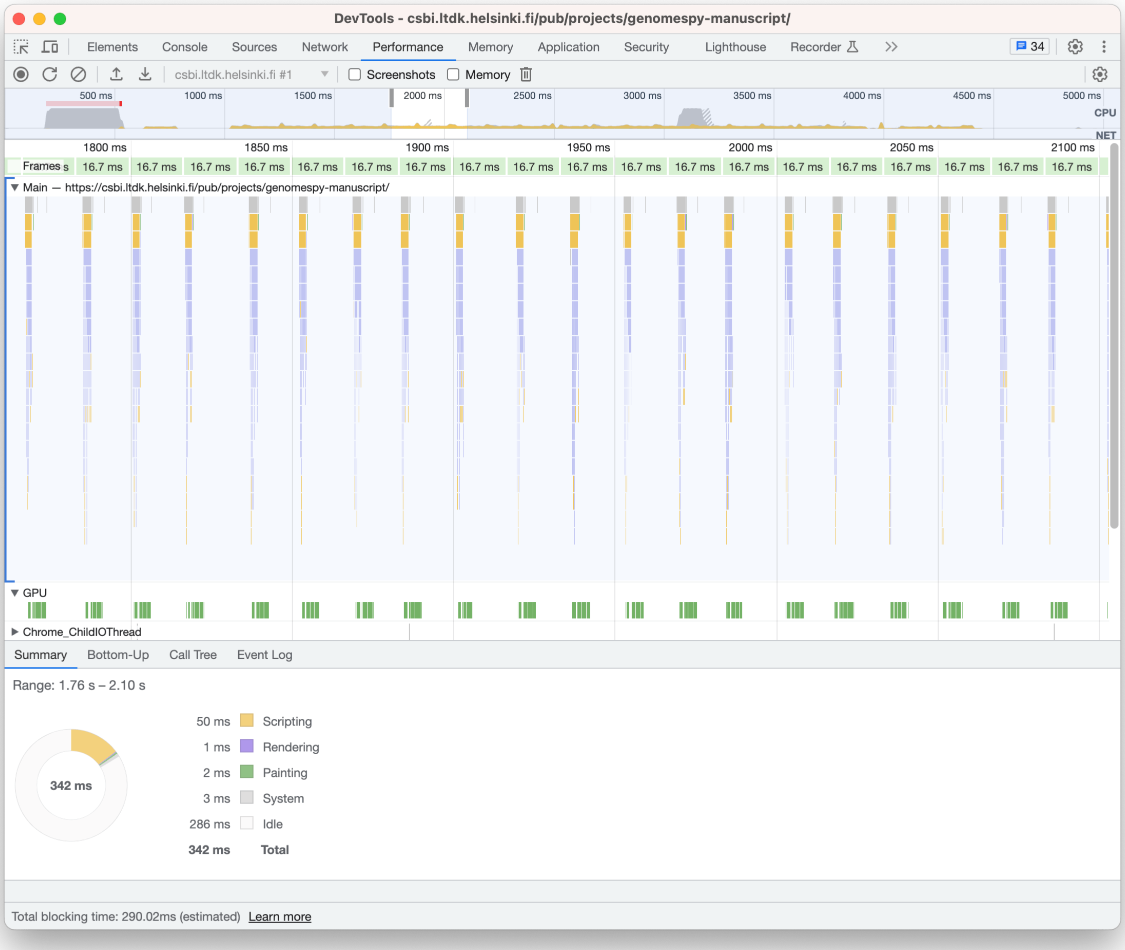

Supplementary Figure 3: Chrome Developer Tools profile of GenomeSpy when zooming the whole dataset (753 samples) in the visualization shown in Figure 1c (https://csbi.ltdk.helsinki.fi/p/genomespy-preprint/). The profile shows very low CPU utilization, which guarantees smooth animation without dropped frames.

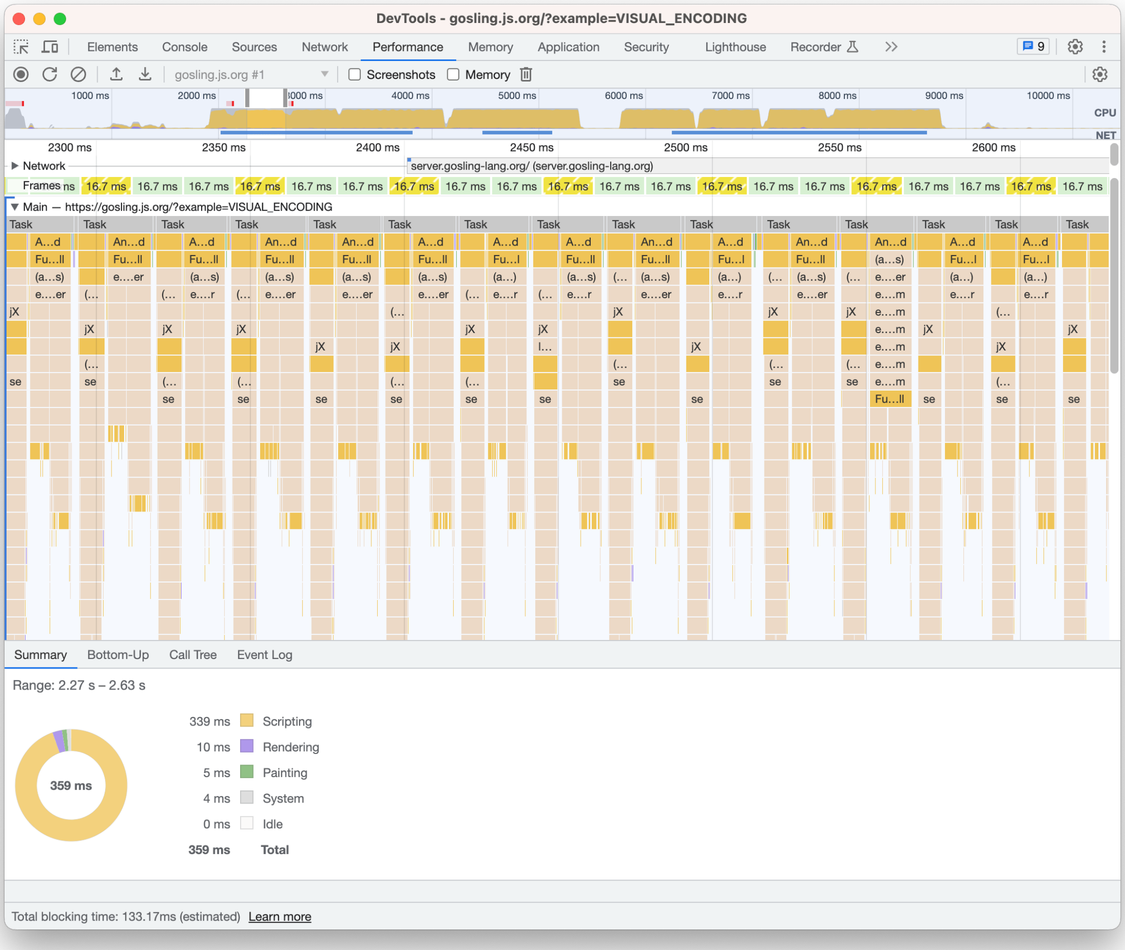

Supplementary Figure 4: Chrome Developer Tools profile of Gosling when zooming the Visual Encoding example (https://gosling.js.org/?example=VISUAL_ENCODING). The profile shows full CPU utilization and multiple dropped frames, resulting in choppy animation during interactions.
